# Sparse gut microbiomes in solitary bees and wasps

**DOI:** 10.64898/2026.07.02.736198

**Authors:** Jen N. Schlauch Saiyawong, Kristal M. Watrous, Stephen L. Buchmann, Annalie Melin, Tobin J. Hammer

**Affiliations:** University of California, Irvine, Department of Ecology and Evolutionary Biology; University of Arizona, Departments of Ecology and Evolutionary Biology, and Entomology; Compton Herbarium, South African National Biodiversity Institute, Claremont, South Africa; African Centre for Coastal Palaeoscience, Nelson Mandela University, Port Elizabeth, South Africa

**Keywords:** microbiota, pollinators, Hymenoptera, Masarinae, sociality, pollen-feeding, Anthophila

## Abstract

Bees and wasps are ecologically vital, but many species are declining due to anthropogenic stressors. Social bees harbour host-specific and dense gut microbiomes that affect their resilience to stress. However, there are tens of thousands of other bee and wasp species that vary in sociality and diet (including pollen-feeding and predatory guilds), traits known to influence host-microbe symbioses. The role of gut microbes in the biology of these species is largely unknown. Here, we measured the composition and absolute abundance of bacterial communities in adult abdomens across 61 genera and 14 families of field-collected bees, predatory wasps, and pollen wasps. We found that solitary bees and both wasp guilds harbor distinct bacterial taxa and lower bacterial abundances as compared with social bees. Bacterial abundances also varied extensively among and within genera of solitary bees, with little variation explained by body size, diet breadth, or nesting ecology. Further, microbiome composition was only weakly differentiated among solitary bees and the two wasp groups, even comparing herbivorous (pollen-feeding) and carnivorous taxa. We suggest that the sparse and somewhat stochastic microbiomes of solitary bees and wasps reflect weak host dependence on microbially mediated functions, a trait that may influence their responses to environmental change.

## Introduction

Bees and wasps are an enormously diverse group with essential roles in terrestrial ecosystems, but their persistence is threatened by anthropogenic stressors [1–4]. How these insects will fare in the future may depend, in part, on the nature of their microbiomes. Many animal taxa harbour gut microbial symbionts that improve their ability to resist diseases, pesticides, nutrient limitation, or other perturbations [5–7]. Others appear to lack beneficial gut symbionts, instead harbouring only transients, commensals, or pathogens [8], and may thus lack an important mechanism for adapting to changing environments [9,10]. How common these distinct symbiotic strategies are among most bees and wasps is unclear. While a few taxa have been thoroughly studied, generalizing findings is complicated by variation in sociality and diet, traits known to shape the gut microbiome [11,12].

Much of the research on gut microbiomes of bees and wasps has focused on eusocial species, particularly honeybees (*Apis*), bumblebees (*Bombus*), and certain wasps in the family Vespidae. Adult *Apis* and *Bombus* host a highly abundant (∼10^8^-10^9^ cells per individual) and consistent microbial community of host-specific gut bacteria that are transmitted between generations within the colony [13–15]. This symbiosis appears to be ancient [14,15], likely reflecting host dependence on bacterially provided functions, including digestion, detoxification, and pathogen defense [16]. Among wasps, one eusocial species with large and long-lived colonies, as well as an unusual diet—the honey wasp *Brachygastra mellifica*—was also found to harbour abundant and host-specific gut bacteria [17].

Over 77% of bee species and the vast majority of wasp species are solitary [1,18,19], and 16S rRNA gene sequencing data indicate that their gut microbiomes differ from those of the aforementioned eusocial species. In adult solitary bees and wasps, gut bacterial communities often vary substantially in composition between conspecific individuals or populations, and their constituent taxa appear to be horizontally acquired from flowers, prey, or other environmental sources [17,20–23]. These characteristics are consistent with a weak dependence on functions provided by bacterial symbionts [8], although counter-examples exist [24]. To our knowledge, experimental data are not available to evaluate the hypothesis of weak gut-symbiotic dependence in adult solitary bees and wasps, as most species cannot be easily reared in the laboratory. Therefore, there is a need for an observational approach to broadly assessing gut symbiosis in these insects.

Here we use absolute abundance—an often overlooked parameter in microbiome studies [25,26]—to infer the functional importance of gut bacterial communities to solitary bees and wasps. Absolute abundance reflects function because a low number of microbes in a microbiome will constrain the total metabolic or ecological outputs of that microbiome. Thus, hosts that rely upon beneficial symbionts typically provide nutrients and housing to support high levels of microbial growth and hence microbial services [27–29]. In contrast, hosts that do not invest in maintaining beneficial symbionts—those with largely transient or commensal microbes—are expected to harbour fewer microbes overall, the abundance of which may fluctuate stochastically depending on diet-derived microbial inputs or other factors [30–32]. The few studies that have quantified gut microbiomes in adult solitary bees or wasps have found low absolute abundances compared with microbiomes in eusocial species [14,17], but much broader sampling is needed to infer whether this is a general pattern.

To assess the role of sociality in shaping the gut microbiome across bees and wasps, we compared bacterial communities among field-collected solitary bees (35 genera), eusocial bees (6 genera), predatory wasps (15 genera, two of which are eusocial), and pollen wasps (3 genera). The latter group, one of only two obligately pollen-feeding Hymenopterans beside bees [33,34], was included to examine the role of diet as an additional factor shaping the microbiome.

Adult-stage abdomens (technically, metasomas) were sampled, which include most of the gut (excluding the proximal esophagus or crop) but also reproductive tissues, hemolymph, and other potential sites of symbiont colonization. We used 16S rRNA gene sequencing and quantitative PCR to measure bacterial community composition and absolute abundance, respectively. Our work reveals large and previously unexplored variation in bacterial abundances among adult bees and wasps, at both individual- and species-level scales. Such variation has the potential to influence host fitness and the resilience of these insects in the Anthropocene.

## Materials and Methods

### Field collections

A total of 443 wild adult bee and wasp specimens were sampled for this study. Of these, 310 specimens were collected in California and Arizona from April to August of 2023 and stored in 95% ethanol. An additional 100 samples from South Africa, the Canary Islands, and the western United States were included; these were collected via hand net between January 2022 and March 2025 and stored in 95% ethanol. Thirty of the South Africa specimens were euthanized via cyanide killing jar, stored at -20 °C briefly, then transferred to 95% ethanol. Thirty-three Meliponini samples, stored in 95% ethanol, were included in the dataset; these samples are described in [35] and were previously sequenced [36]. All specimens besides the Meliponini were identified at least to genus by J. Schlauch using keys [37,38].

### Bee trait data

We measured the body size of solitary bees and a subset of eusocial bees using a 5mm Mini Measuring Ruler and dissection microscope. Intertegular distance (ITD), a standard metric of body size for bees, was measured between the tegulae at the wing bases [39]. Nesting ecology of solitary bees was summarized from [1] and classified as: ground nesting, leaf-cutter, stem-nest, and brood parasite. Diet breadth of solitary bees was classified as generalist or specialist [1]. Sample metadata are provided through Dryad (DOI: 10.5061/dryad.4qrfj6qt2).

### DNA sequencing and qPCR

DNA was extracted from individual homogenized abdomens and PCR-amplified with V4 (515F/806R) 16S rRNA gene primers. Following established recommendations for controlling contamination [40], multiple blanks (negative controls) and mock communities (positive controls) were included. Amplicons were sequenced on four Illumina Miseq runs (2 × 250 bp or 400 bp, depending on the run). Raw reads are available on NCBI (BioProject #PRJNA1472269). Stingless bee samples from ref. [41] were added to this dataset; these were originally extracted and PCR-amplified using the same protocol in the same laboratory, with 16S rRNA V4 amplicons sequenced on an Illumina MiSeq (2 × 250 bp). All sequence data were quality-filtered and processed with DADA2 [42] following ref. [43]. The ASV table and representative sequences are available through Dryad (DOI: 10.5061/dryad.4qrfj6qt2). Quantitative PCR (qPCR) with 27F/355R primers was used to measure absolute abundances of bacteria in the DNA extracts, following refs. [42,47]. We multiplied the resulting total 16S copy numbers by the proportion of V4 amplicon sequences classified as non-contaminant bacteria, producing an estimate of bacterial 16S rRNA gene copies per abdomen. More details are provided in the Supplemental Methods.

### Statistical analysis

Analyses were conducted in R v. 4.3.2 (R Core Team, 2023). To test whether chloroplast sequences or absolute abundances varied between host guilds, linear mixed-effects models were fitted with either chloroplast proportion or absolute abundance (log-transformed) as the response, host guild as a fixed effect, and host genus as a random effect. The chloroplast model used a likelihood ratio test, with proportion comparisons based on Tukey’s HSD. To test whether diet, nesting ecology, or body size (intertegular distance) was associated with absolute abundance, identical models were fitted for solitary bees only, with diet or nesting ecology as the fixed effect [44,45]. To visualize microbiome composition across host guilds, we used an NMDS ordination of Bray–Curtis dissimilarities with vegan [46]. PERMANOVA was performed to test for differences among guilds in composition. Pairwise comparisons were conducted using pairwiseAdonis [47] with False Discovery Rate correction. See the Supplemental Methods for further details.

## Results & Discussion

After sequence data processing, we obtained microbiome composition (16S rRNA gene sequencing) and absolute abundance (qPCR) data from 422 adult bee and wasp specimens, including 298 solitary bees from 35 genera in 6 families; 66 eusocial bees from 8 genera in 1 family; 36 predatory wasps from 15 genera in 6 families; and 22 pollen wasps from 3 genera in 1 family. This dataset likely represents the broadest (in terms of phylogenetic extent) gut microbiome characterization of bees and wasps investigated to date. Further, the host taxa differ in sociality and diet (Fig. 1A), allowing an assessment of how these traits may be linked to the gut microbiome.

**Figure 1.**
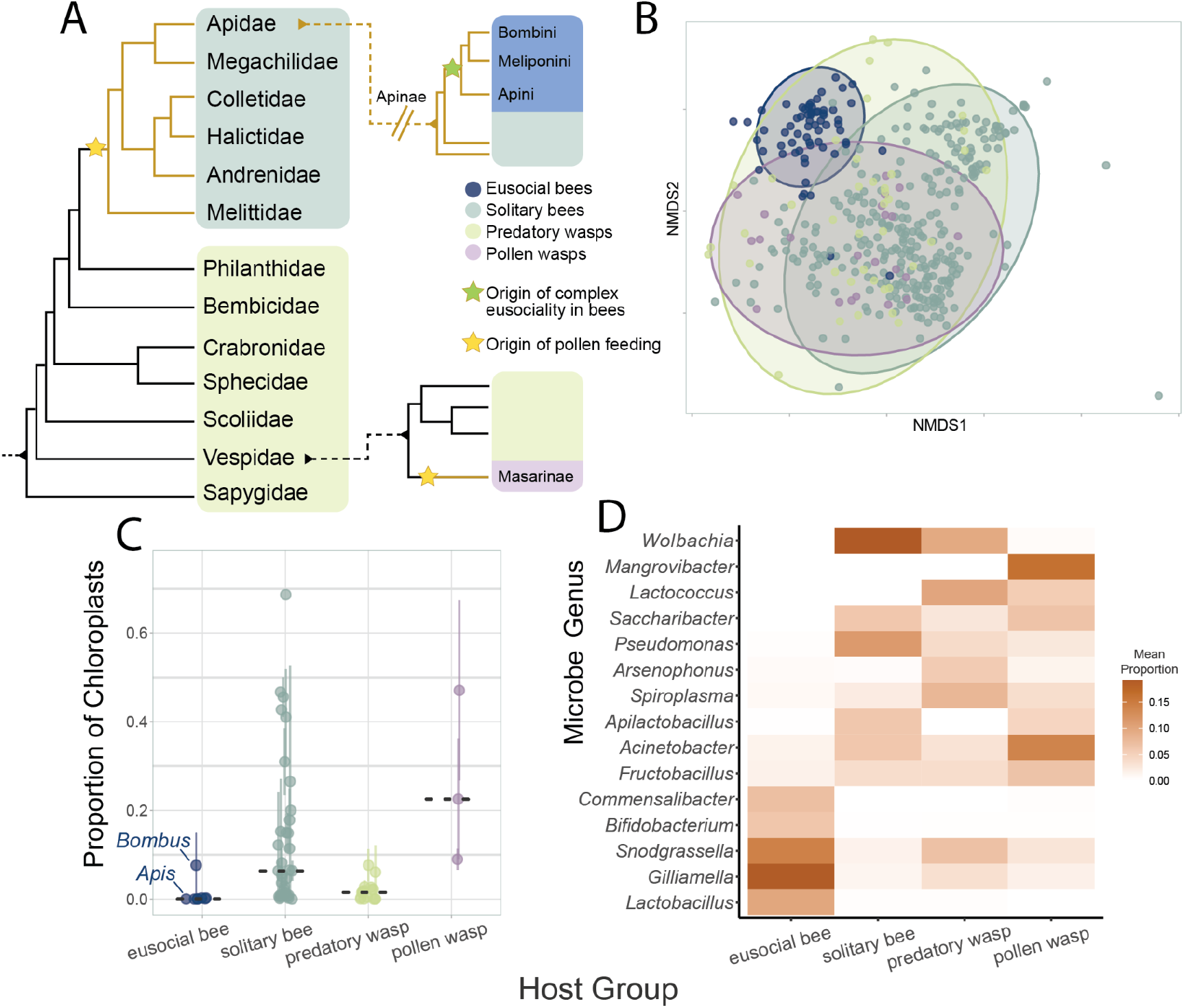
Variation in microbiome composition across adult bees and wasps. (A) Cladogram of all families included in the study, adapted from [81], indicating the origin of complex eusociality within bees and the independent origins of pollen-feeding in bees and pollen wasps. (B) NMDS ordination of Bray-Curtis dissimilarities. Each point is an individual sample. Ellipses represent 95% confidence intervals around the centroid for each insect guild. (C) Chloroplast content in sequence libraries. Each point represents the mean proportion of chloroplast sequences for a given host genus; for genera with replicate individuals, the vertical line indicates the SEM. Medians for each host guild are indicated by dashed grey lines. Points are horizontally jittered. Among eusocial bees, honeybees (Apis) and bumblebees (Bombus) are labeled, but not stingless bee (Meliponini) genera. (D) Fifteen of the most common bacteria in bees and wasps, representing a combination of, for each host guild, the five bacterial genera with the highest mean proportion of sequences across samples.

Microbiome composition varied between eusocial bees, solitary bees, predatory wasps, and pollen wasps (“host guilds”) (PERMANOVA; F = 17.622, R^2^= 0.11, p = 0.001; Fig. 1B). All host guilds differed in microbiome composition from one another (pairwise PERMANOVA; p-adjusted = 0.001), but eusocial bees were more different to the solitary bees and wasps (R^2^ values all > 0.11) than solitary bees and wasps were to each other (R^2^ values all < 0.04). This result matches previous evidence that eusocial bees have a highly distinctive gut microbiome relative to solitary bees [14,12]. In contrast, there was relatively little partitioning by diet, with large overlap in composition among predatory wasps and the herbivorous solitary bees and pollen wasps (Fig. 1B,D). This is unusual, because in other animals, carnivores and herbivores typically have very different gut microbiome composition [11,48].

The overlap in microbiome composition between solitary bees and the two wasp guilds might be driven by horizontal transmission from common pools of environmental microbes. Most of the prevalent bacterial genera in solitary bees and pollen wasps (e.g., *Pseudomonas, Acinetobacter, Apilactobacillus, Saccharibacter, Fructobacillus, Lactococcus*; Fig. 1D) frequently inhabit flowers [21,49,50]. These results align with prior research demonstrating that adult solitary bees acquire gut bacteria from flowers or other environmental substrates [51,52]. Some of these putatively flower-derived bacteria (e.g., *Lactococcus* and *Fructobacillus*) were also common in predatory wasps (Fig. 1D), potentially because they often visit flowers to consume nectar or prey on florivorous arthropods [19]. *Wolbachia*, a vertically transmitted endosymbiont that frequently parasitizes insect reproductive systems [53], was also very common in solitary bees and predatory wasps (Fig. 1D), as expected from prior surveys of Hymenoptera [54,55]. The eusocial bee microbiomes were dominated by gut bacterial taxa generally thought to be specific to eusocial bees [14,15], especially *Snodgrassella* and *Gilliamella* (Fig. 1D). However, these bacteria also occurred at a low prevalence in other bee and wasp samples (Fig. 1D). One possibility is that they are not as eusocial-bee-restricted as previously thought [56]. Another explanation is cross-contamination among samples during DNA extraction, library preparation and sequencing, an issue exacerbated by low absolute abundance of microbes[40]. For the predatory wasps in particular, eusocial bee-specific bacteria could also be present due to recent handling or consumption of bee prey [23].

Off-target sequences such as chloroplasts can provide clues into gut microbial abundance in herbivorous animals [40]. We found that the proportion of chloroplast sequences varied between host guilds (linear mixed effects model; χ^2^= 15.24, p= 0.002). As expected from their diet, predatory wasps have few chloroplast sequences (Fig. 1C). Among the remaining groups, all of which ingest pollen as adults [1,38,57], solitary bees and pollen wasps have higher proportions of chloroplast sequences than eusocial bees, although the difference was only statistically significant with pollen wasps (pairwise post hoc Tukey test; p = 0.07 and p = 0.04, respectively) (Fig. 1C). In other herbivorous insects, high levels of chloroplasts in sequencing data reflect a low absolute abundance of gut bacteria [30,58].

We used qPCR to directly measure the absolute abundance of bacteria (in terms of 16S rRNA gene copies) and combined it with the sequencing data to correct for nonbacterial amplification. Bacterial abundance varied strongly among host guilds (linear mixed effects model; χ^2^= 27.312, p < 0.001), with solitary bees, predatory wasps, and pollen wasps harbouring, on average, orders of magnitude (∼2000X) fewer bacteria than eusocial bees (Fig. 2A). The difference in absolute abundance between eusocial bees and the other host guilds would be larger if only honeybees and bumblebees were compared; as observed previously [14], some stingless bee genera have much less gut bacterial biomass (Fig. 2A). Further, the difference in bacterial abundances would also likely be larger if only gut-associated bacteria were compared. *Wolbachia*, the most common bacterium in the abdomens of solitary bees and second-most common in predatory wasps (Fig. 1D), inhabits reproductive tissues rather than the gut (Werren et al. 1995). In contrast, all dominant bacterial genera in eusocial bee abdomens (Fig. 1D) have been found to specifically inhabit the gut [12,14]. A large discrepancy in gut bacterial biomass between solitary bees and eusocial bees, as indicated here by qPCR, is also supported by scanning electron microscopy in the solitary bee *Megachile tosticauda* and honeybees [59].

**Figure 2.**
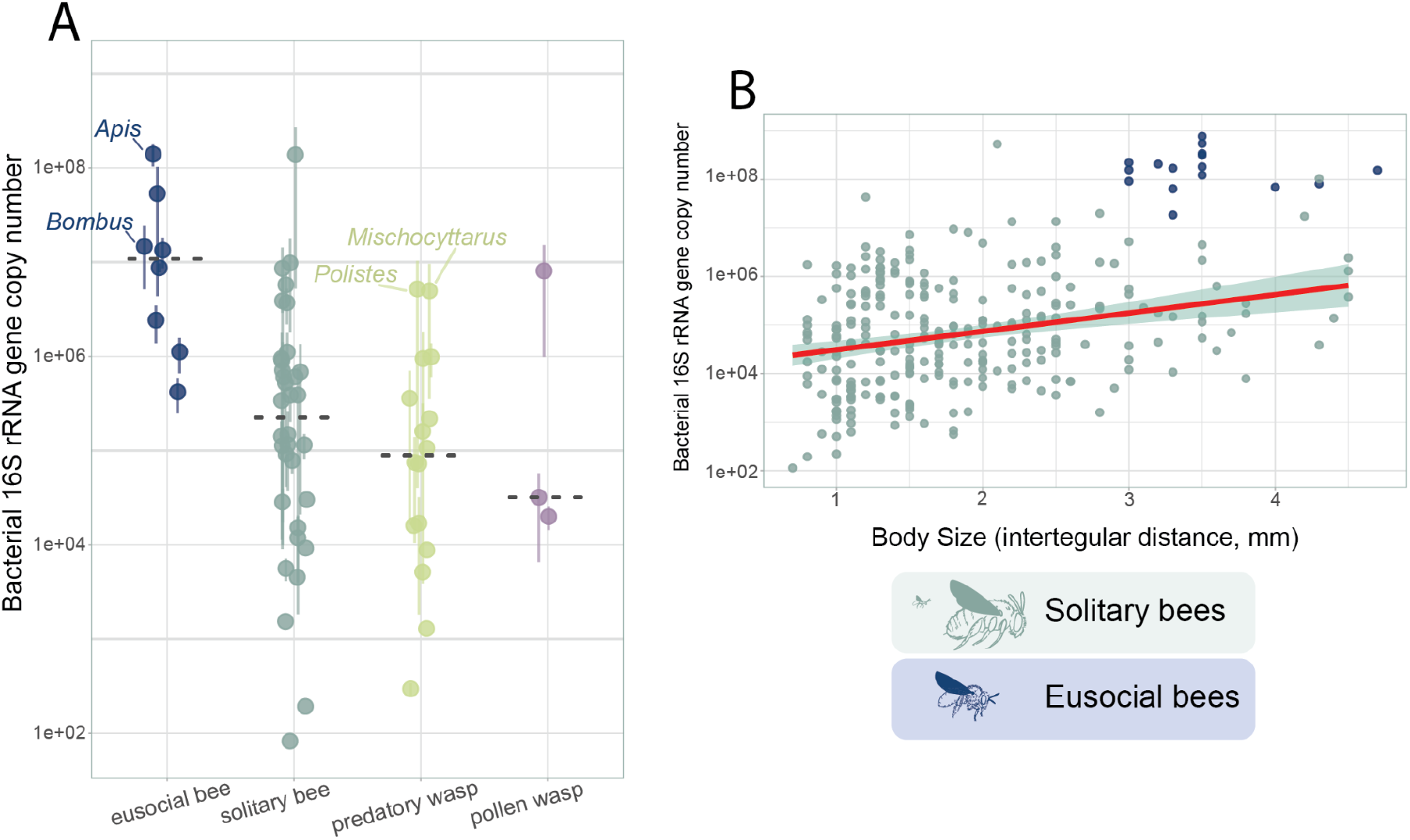
Variation in absolute abundances of bacteria across bees and wasps. (A) Bacterial 16S rRNA gene copy numbers per adult abdomen across bee and wasp guilds. The data and sample sizes are presented as in Fig. 1C. (B) Association between body size and bacterial 16S rRNA gene copy number for solitary bees only (N = 277 specimens). The red line represents a linear regression fit to the solitary bee data. A subset of eusocial bee specimens (N = 15 Apis, 1 Bombus) is included for reference to illustrate their body size to copy number relationship, but these data are not included in the regression model.

Solitary bees, predatory wasps, and pollen wasps also exhibit orders-of-magnitude levels of variability in bacterial abundance, both within and among genera (Fig. 2A). We examined whether such variability could be explained by host traits, focusing on solitary bees, for which we had the largest sample size and the most trait information. At the level of individual specimens, solitary bee body size is positively, but only weakly associated with bacterial abundance (linear mixed effects model; χ^2^= 17.46, p < 0.001, R^2^ = 0.080, Fig. 2B). At the level of genera, neither diet breadth (linear mixed-effects model, χ^2^=0.7527, p = 0.7627) nor nesting ecology (linear mixed effects model; χ^2^= 2.5645, p = 0.5573) are associated with bacterial abundance. One possible driver of this apparently stochastic variation is that individuals with especially high bacterial abundances were infected by opportunistic pathogens or parasites. Alternatively, if most of the microbiome is acquired from the environment, as suggested above, and its growth is not tightly regulated by the host, then an individual bee or wasp’s microbial load may simply reflect the microbial load of its most recent meal [30].

We argue that the low and variable bacterial abundances observed in solitary bees, predatory wasps, and pollen wasps likely reflect a generally weaker dependence on gut symbionts as compared with eusocial bees. Why might eusociality affect symbiotic dependence? One possibility is that challenges of living in large groups, such as greater disease risk or nutritional demands, increase the selective pressure to maintain beneficial symbionts that help hosts meet those challenges [36]. Overlap of adult generations, a defining characteristic of eusociality [60], also provides a ready-made route for intergenerational transmission of microbes, in turn enabling the kind of persistent, reciprocal interaction that is necessary for coevolution [14,61].

As only two predatory wasp genera in our sample set, *Polistes* and *Mischocyttarus*, are eusocial, we cannot conclusively test the effect of sociality on gut microbiome abundance within wasps. However, we note that the absolute abundances of bacteria in these two genera were higher than all other predatory wasp taxa (Fig. 2A). A previous study found that eusocial, predatory *Polistes* wasps had gut bacterial abundances slightly higher than solitary wasps, but still much lower than eusocial honey wasps [17]. Among ants, all of which are eusocial, many omnivorous species have low-abundance gut microbiomes [26,62]. At least among Hymenoptera, perhaps evolution of both specialized diets and eusociality are necessary to promote host dependence on (and investment in) beneficial gut bacterial symbionts.

That an animal could exist without large numbers of beneficial gut symbionts may be particularly surprising for herbivorous insects like most solitary bees and pollen wasps, given that herbivory often fundamentally requires microbes to supply digestive enzymes or missing nutrients [63]. However, endogenous (host-encoded) biochemical mechanisms for pollen digestion have been described in bees [64–66], and nectar and pollen are considered to be a nutritionally complete resource for insects [67]. Indeed, sterile honeybees and bumblebees have been shown to exhibit similar survival rates as those harbouring their core gut bacterial symbionts [68,69], while a lack of pollen nutrients does decrease adult survival [70]. This indicates that a gut microbiome is not strictly required for pollen digestion or adequate nutrition.

A weak dependence on a gut microbiome, whether for pollen-feeding or other functions, could influence the resilience of solitary bees and wasps in a changing environment. On the one hand, a lack of abundant, beneficial gut symbionts may mean that these insects are missing a source of plasticity that could help them deal with changing environments [71,72]. For example, eusocial bee gut microbiomes provide pathogen resistance [5,73], and pathogens are a major contributor to bee declines [74]. On the other hand, solitary bees and wasps may be less affected by certain stressors than strongly microbiome-dependent insects. Microbial symbionts can have physiologies that are uniquely sensitive to particular stressors, acting as an “Achilles’ heel” for their host [75]. For example, heat-sensitive gut symbionts constrain the thermal tolerance of stinkbugs [76], and some of the core honeybee gut symbionts are inhibited by the herbicide glyphosate [69]. Manipulative experiments will likely be needed to determine which of these scenarios applies to solitary bees and wasps, and with which kinds of stressors.

There are important caveats to our findings. For example, our analyses were limited to adults, because immature stages are much more difficult to sample from a broad range of species. In some bee taxa (including honeybees), developing larvae appear to have a low-abundance and/or diet-derived microbiome [12,22,77,78]. Thus, in some solitary bees and wasps, it is possible that neither larvae nor adults strongly depend on gut symbionts. But this does not necessarily apply to all species or life stages [79,80]. Further, bees and wasps are phylogenetically and ecologically diverse, and our sample set represents only a small fraction of this biodiversity. Nevertheless, our findings provide a novel null hypothesis for future research on solitary bees and wasps: that these insects generally harbour sparse gut microbiomes that provide minimal benefits to their host.

## Acknowledgements

We thank M. J. Larson and M. Huang for their help with specimen processing, identification assistance from J. Palewek, and collectors who contributed specimens: C. Rasmussen, A. S. Nelson, L. Pettway, H. Mahoney, E. Muramoto, A. Vazquez, J. Cowles, Logan Bee Lab. J.N.S. was supported by the U.S. National Science Foundation Graduate Research Fellowship (Award #DGE-1839285), and grants from the Sea and Sage Audubon Society, California Native Plant Society Orange County Chapter, and University of California Nature. A.M. received postdoctoral funding from the Mapula Trust. We are grateful for fieldwork support from several UC Nature Reserves, Pogo Hill, Mojave Desert Land Trust, Tijuana Slough NWR, IRWD San Joaquin Marsh, and the Desert Studies Center.

## Supplemental Methods

### DNA Extraction and Sequencing

After taxonomic identification, abdomens were removed and placed in individual tubes containing 800μL ZymoBIOMICS lysis solution (Zymo Research), homogenized with a pestle, then added to ∼0.5mL autoclaved zirconia silica beads (BioSpec) and two 5mm 440C stainless steel grinding balls. Samples were homogenized for six minutes with 1 minute rests every 2 minutes using a Benchmark BeadBlaster 96 homogenizer. Sample homogenates were then stored at -70ºC until extraction of genomic DNA (gDNA) with the ZymoBIOMICS DNA 96 kit. Following best practices for controlling contamination during gDNA extraction and library preparation [40], 28 negative controls (blanks) and 13 replicates of the ZymoBIOMICS Microbial Community Standard (Zymo Research) were included and sequenced alongside the samples. gDNA was PCR-amplified using barcoded universal primers (515F/806R) targeting the V4 region of the 16S rRNA gene, following the V.1 protocol used by the Earth Microbiome Project [82]. Clean-up and normalization were performed using the SequalPrep kit (Applied Biosystems). Pooled amplicons were run on four separate sequencing runs: two Illumina MiSeq (2 × 400 bp paired end reads) runs at the University of California Irvine Genomics Research and Technology Hub (UCI GRTH), one Illumina MiSeq (2 × 250 bp paired end reads) at UCI GRTH, and one Illumina MiSeq (2 × 250 bp) run at the University of Colorado Boulder CIRES Microbial Community Sequencing Lab. Raw reads are available on NCBI (BioProject #PRJNA1472269)

### Sequence data processing

Raw sequences were processed using a previously described protocol [45]. Briefly, DADA2 [42]was used to trim and quality-filter reads (maxEE = 2, truncQ = 2, truncLen = 240 fwd and 200 rev), infer amplicon sequence variants (ASVs), and remove chimeric ASVs. The RDP Naive Bayesian Classifier [83] was used to assign taxonomy to the ASV table, with the SILVA 138.2 SSU rRNA reference database [84]. The ASV table and representative sequences are available through Dryad (DOI: 10.5061/dryad.4qrfj6qt2).

The ASV table was loaded into R version 4.3.2 and further analyzed using the mctoolsr R package (https://github.com/leffj/mctoolsr). Samples with zero reads were dropped, leaving 481 samples. We filtered out ASVs with fewer than 10 reads across all samples (5,587 removed), ASVs without a classification to the domain level (18 removed), as well as mitochondria, chloroplasts, and other eukaryotic ASVs (1,114 ASVs removed). We used data from the mock communities to validate the accuracy of our sequencing. The top ten ASVs, in order of mean relative abundance across the 13 samples, were: *Escherichia-Shigella, Bacillus, Salmonella, Pseudomonas, Limosilactobacillus, Listeria, Salmonella, Enterococcus, Staphylococcus*, and *Candidatus Schmidhempelia*. All of these taxa are in the mock community except for *Schmidhempelia*, a bumblebee-specialized bacterium, which probably occurred in the mock communities due to cross-contamination from bumblebee samples during DNA extraction. Contaminant ASVs were identified using the isContaminant function in the decontam package [85] with the “prevalence” method, resulting in 91 out of 8992 ASVs classified as contaminants, which were then removed from the dataset. We then curated the taxonomy of the ASVs to avoid lumping different taxa with “NA” as genus by merging their “unid” names with the order and family names. The mean library size of abdomen samples was 23,356; to maximize the number of samples available for analysis, samples were rarefied (randomly subsampled) to 500 sequences each, resulting in 422 samples (12 removed). Because the libraries originated from multiple sequencing runs with different read lengths, we summarized the ASV table to the genus level for further analyses.

### Absolute abundance

Quantitative PCR (qPCR) was performed to measure 16S rRNA gene copy numbers. We used universal 27F/355R primers targeting the V1-V2 region of the 16S rRNA gene, following a previously described protocol [43], and a qTOWER3 system (Analytik Jena). Reactions were run in 10μL volumes in triplicate, using PowerUp SYBR Green Master Mix (ThermoFisher) and 1μL of template DNA per reaction. Nine negative control samples (two PCR no-template controls and seven extraction blanks) were included; the mean copy number (16.94 copies/uL) was subtracted from the abdomen samples. To calculate total 16S copy numbers per abdomen, the mean copy number per uL across triplicates was multiplied by 20 (as gDNA was eluted in 20uL) and by 4 (as only ¼ of each abdomen homogenate was used in the extraction). Finally, for each sample, the latter value was multiplied by the proportion of amplicon sequences classified as non-contaminant bacteria, producing an estimate of the number of bacterial 16S rRNA gene copies per abdomen.

### Statistical Analyses

Statistical analyses were conducted in R v. 4.3.2 (R Core Team, 2023). Linear mixed-effects models were fitted using the lmer() function in lme4 [44]. We tested the significance of fixed effects with Wald χ^2^ tests with type III sums of squares (car::Anova [86]), except for the chloroplast models, which used likelihood ratio tests (lme4::anova, Bates et al. 2015). Comparisons of chloroplast proportions were based on pairwise post hoc Tukey’s HSD tests, using the effect sizes of the estimated marginal means of host guilds (emmeans::pairs [45]). To test whether chloroplast sequences or absolute abundances varied between host guilds, linear mixed-effects models were fitted with either chloroplast proportion or absolute abundance (log-transformed) as the response, host guild as a fixed effect, and host genus as a random effect. To test whether diet or nesting ecology was associated with absolute abundance, identical linear mixed-effects models were fitted for solitary bees only, with diet or nesting ecology as the singular fixed effect and host genus as a random effect. To test the effect of bee body size (ITD) on absolute abundance, a linear model was generated for solitary bees only with absolute abundance (log-transformed) as the response and ITD as the predictor.

To visualize microbiome composition across host guilds, we used non-metric multidimensional scaling ordination (NMDS) of Bray–Curtis dissimilarities calculated with the vegan package [46]. A permutational analysis of variance (PERMANOVA) with 999 permutations was performed to test for differences among guilds in composition. Pairwise comparisons were conducted using pairwiseAdonis [47] with False Discovery Rate correction.

